# Sex differences in the effects of mild traumatic brain injury and progesterone treatment on anxiety-like behavior and fear conditioning in rats

**DOI:** 10.1101/2023.01.13.523944

**Authors:** Laura C. Fox, Jamie L. Scholl, Geralyn M. Palmer, Gina L. Forster, Michael J. Watt

## Abstract

Mild traumatic brain injuries (mild TBIs) commonly occur in young adults of both sexes, oftentimes in high-stress environments. In humans, sex differences have been observed in the development of post-concussive anxiety and PTSD-like behaviors. Progesterone, a sex steroid that has neuroprotective properties, restores cognitive function in animal models following more severe TBI, but its effectiveness in preventing the psychological symptoms associated with mild TBI has not been evaluated. Using a model of mild TBI that pairs a social stressor (social defeat) with weight drop, male and naturally estrous-cycling female rats were treated with 4 mg/kg progesterone or vehicle once daily for 5 days after injury. Behavioral measures, including elevated plus maze (EPM), contextual fear conditioning, and novel object recognition (NOR) were assessed following progesterone treatment. Anxiety-like behavior was increased by mild TBI in male rats, with a smaller effect seen in female rats in the diestrus phase at the time of EPM testing. In contrast, mild TBI impaired fear learning in female rats in estrus at the time of fear acquisition. Progesterone treatment failed to attenuate post-mild TBI anxiety-like behavior in either sex. Furthermore, progesterone increased fear conditioning and impaired NOR discrimination in male rats, independent of TBI status. Overall, both sex and estrous cycle contributed to psychological outcomes following mild TBI, which were not ameliorated by post-TBI progesterone. This suggests sex steroids play an important role as a moderator of the expression of mild TBI-induced psychological symptoms, rather than as a potential treatment for their underlying etiology.

## Introduction

Over 85% of the 2.5 million traumatic brain injuries (TBIs) diagnosed in the United States each year are classified as mild (CDC 2015), which entail no penetrative head wound and result in loss of consciousness for less than 30 minutes (Brasure et al., 2012). Also called concussions, these injuries frequently happen in environments involving sport, warfare, or domestic violence (Moore et al., 2006). While post concussive symptoms such as cognitive impairment, mood disruptions, and sleep disturbances appear to resolve spontaneously within 3 months post mild TBI for many patients, a variable subset (15-50%) experience symptoms that may continue months to years beyond the initial impact (Bazarian et al., 2010, Moore et al., 2006; Polinder et al., 2018).

One of the most commonly reported persistent psychological issues following mild TBI relates to anxiety disorders (Kennedy et al., 2007, Moore et al., 2006). For example, 43.9% of returning soldiers who report experiencing a mild TBI also express co-morbid symptoms of posttraumatic stress disorder (PTSD) (Hoge et al., 2008). Symptoms of mild TBI and PTSD often overlap (Hoge et al., 2008, Kennedy et al., 2007, Davies et al., 2016), and these, along with other anxiety behaviors, can arise months after the initial trauma. Although findings are mixed, the risk of developing an anxiety disorder within the 12-48 months post mTBI appears to be higher in patients without any pre-existing affective disorder (Delmonico et al., 2021; Sheldrake et al., 2022), suggesting a causal relationship between mild TBI and anxiety. Damage to cells in limbic structures such as the hippocampus and amygdala have been shown to promote anxiety-like behavior in rodent models of mild TBI (Meyer et al., 2012), and there is some evidence that neuroinflammation also plays a role in emergence of mood and anxiety disorders in humans (Risbrough et al., 2022). However, no effective clinical biomarkers currently exist for reliably determining which individuals will experience long-term anxiety after a mild TBI (Jeter et al., 2013; Polinder et al., 2018), limiting the options for targeting underlying neural mechanisms. In addition, current post-injury therapies for mild TBI-associated anxiety and PSTD-like symptoms are often ineffective (Kennedy et al., 2007). Therefore, an ideal treatment would comprise a prophylactic therapy that prevents neuronal changes associated with increased anxiety, but that does not disrupt function in the >50% of concussive patients who do not develop long-term symptoms (Moore et al., 2006).

Men and women may respond dissimilarly to mild TBI. Although men are more likely to receive a mild TBI, a higher percentage of women report experiencing long-term complications (Bazarian et al., 2010, Styrke et al., 2013). Additional evidence suggests that reproductive cycle phase may play a role in the expression of symptoms. Women who received a TBI while using a hormonal form of oral contraception experienced fewer post-concussion symptoms and negative neurological outcomes than those who were naturally cycling (Wunderle et al., 2014). Ovarian sex steroids, such as estradiol and progesterone, are known to influence anxiety and PTSD symptoms (Albert et al., 2015, Nillni et al., 2015), and have also been shown to possess restorative properties. For example, estradiol has shown some promise in restoring motor and cognitive function following a stroke (Broughton et al., 2014, Waddell et al., 2016), and progesterone has anti-apoptotic (Zhang et al., 2016) and neuroprotective effects (Yousuf et al., 2016, Sayeed & Stein, 2009). These characteristics suggest ovarian sex steroids have potential as therapeutic treatments to prevent the neuronal damage that occurs during mild TBI.

Progesterone has been explored as a treatment for the cognitive deficits seen in more severe TBI. Delivery of high-dose progesterone (4 mg/kg) for 3 or 5 consecutive days shortly after a moderate-to-severe TBI improved cognitive performance in both male and female rats, as assessed by spatial learning ability in a Morris Water Maze (Shear et al., 2002). Progesterone at the same dose also reduces cerebral edema after moderate-to-severe TBI in rats, even when given a day after injury (Roof et al., 1996), and protects against lipid peroxidation that can promote increased vasculature damage and ischemia following severe TBI (Hoffman et al., 1997; Roof et al., 1997; Shear et al., 2002). Further, this dose of progesterone appears to be well tolerated and effective in both sexes (Roof et al., 1992, 1996; Shear et al., 2002). Taken together, these studies highlight the ability of progesterone to aid in preserving cognitive function after severe head trauma. However, whether progesterone alleviates anxiety- and PTSD-like symptoms associated with mild TBI has not been investigated.

This study aimed to first investigate whether the sex differences reported in humans in response to mild TBI could be modeled in both male and naturally estrous-cycling female rats, and second, to understand if progesterone could be beneficial post-injury in preventing emergence of anxiety- and PTSD-like behaviors. A closed-head model of concussive injury concurrent with a social stressor (social defeat) was chosen to stimulate the stress and arousal condition of the environments in which most mild TBIs occur, such as in interpersonal violence, sport, or military conflict (Hoge et al., 2008, Davies et al., 2016; Fox et al., 2016). This model reliably produces increased anxiety-like behavior, enhanced fear conditioning and poorer extinction without producing gross motor deficits (Meyer et al., 2012; Davies et al., 2016; Fox et al., 2016), allowing potential prophylactic effects of progesterone in both male and female rats to be determined effectively.

## Materials and Methods

### Animals

All experiments were approved by the Institutional Animal Care and Use Committee of the University of South Dakota and were carried out in accordance with the National Institutes of Health *Guide for the Care and Use of Laboratory Animals*, and all efforts were made to minimize both the number of animals utilized and potential suffering. Fifty-eight young adult male and 164 naturally estrous-cycling young adult female Sprague-Dawley rats (8 weeks old) were housed in same-sex pairs on a reverse light cycle (12L:12D, lights off 10 AM) with food and water available *ad libitum*. All behavioral testing was performed under red lighting and commenced no earlier than one hour after the dark cycle began. An outline of the experimental timeline is presented in Figure 1. Rats were randomly assigned to one of the following four treatment groups: sham/vehicle, sham/progesterone, mild TBI/vehicle and mild TBI/progesterone.

**Figure 1.**
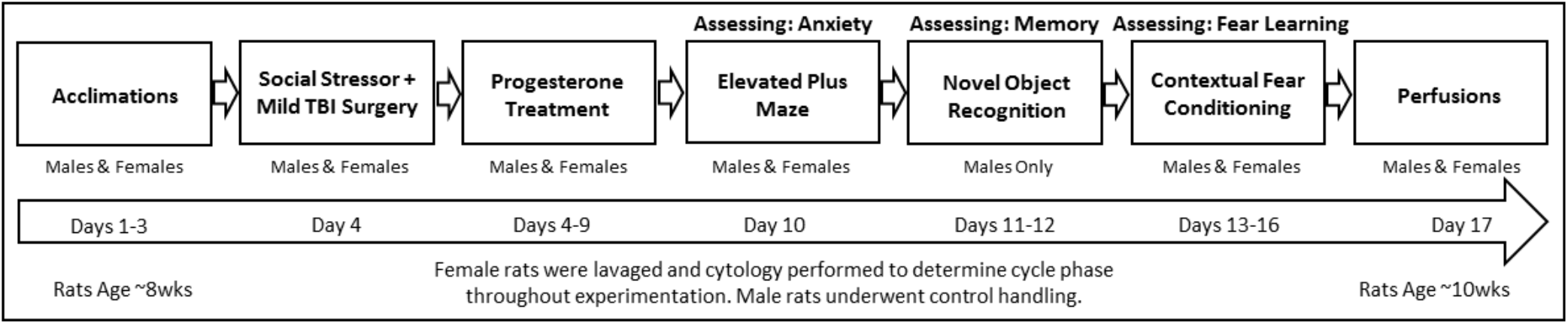
Overview of Methods. Timeline for each experimental animal.

### Estrous Cycle Determination and Vaginal Lavage

Female rats underwent daily lavage beginning at 8 weeks of age, to access one complete reproductive cycle prior to the beginning of experimentation. Similar to methods outlined in Marcondes et al. (2002), approximately 1 mL of filtered 0.9% saline solution was used to flush the female’s vaginal cavity, and the resulting sample was spread across a slide and examined under a microscope at 10 x power. Cytology determined if the female was in metestrus, diestrus, proestrus, or estrus phase (Marcondes et al., 2002). Estrous phase was determined daily throughout the experiment, up to and including the experimental endpoint (perfusion, Fig. 1). Induction of mild TBI had no effect on estrous cycle duration over the duration of the study (sham surgery: 4.760 ± 0.134 days; mild TBI: 4.531 ± 0.071 days; p > 0.1). Male rats underwent daily control handling of equivalent duration. Lavage or male control handling was always performed after behavioral testing, to prevent handling stress from influencing anxiety-like behavior or learning.

### Social Defeat Stressor and Mild Traumatic Brain Injury Induction

Four days after lavage began, all rats were acclimated to a solitary cage and to the testing room (in preparation for social defeat or control conditions) for 40 min/day for three days prior to the treatment day (Davies et al., 2016). On the day of surgery (Fig. 1), rats receiving a mild TBI underwent a single episode of social defeat by a larger, aggressive resident rat of the same sex (>100 g larger than subject), which has been shown to reliably elevate plasma corticosterone in experimental subjects (Davies et al., 2016). Resident males and females were housed in isolation for at least 3 weeks prior to the experiment, to encourage territoriality and aggression (Watt et al., 2014, Davies et al., 2016). After 10 min of physical interaction, subjects remained within the resident’s cage for 30 min, but were confined behind a wire mesh barrier that prevented further physical contact but allowed for transmission of auditory, olfactory and visual cues (Watt et al., 2009; Davies et al., 2016). Defeat episodes were scored on a 1 to 5 scale based on a combination of the number of submissions, attacks, and the overall intensity of the defeat. Control subjects were placed in an empty cage similar to acclimation days. Surgery to induce mild TBI was performed immediately thereafter, with all rats that experienced social defeat also receiving mild TBI, while non-defeated controls underwent sham surgery, following the social defeat model of mild TBI validated in Davies et al. (2016).

Details of the surgical procedure to induce mild TBI are given in Meyer et al. (2012). Briefly, at the end of the 40 min defeat / control episode, rats were anesthetized with isoflurane (3-4% in 3.0 L/min O2), the skin on the top of the head shaved, disinfected, and a 2.5 cm long incision made to expose the skull. A calibrated weight-drop device (following Henninger et al., 2005) was used to drop a 175 g weight from a 42 cm height to deliver a 5,477 Newtons/m2 impact to the exposed skull of the rat, with the force distributed across a 10 mm diameter area via a vertical transducer rod positioned just posterior to bregma and centered over the intraparietal suture. Successful induction of mild TBI was confirmed via delayed righting-reflex time post-surgery, and by the absence of more severe damage (skull fracture, major brain or vascular damage) at the conclusion of behavioral testing (Meyer et al., 2012). One female rat was excluded from the study due to skull fracture. Control (sham) animals underwent surgery without receiving an impact.

### Progesterone Treatment

Three hours following surgery, animals received the first of 6 injections of either vehicle (propylene glycol) or progesterone (>99% progesterone, purified from rats, Sigma-Aldrich Co. St. Louis, MO, USA) at a dose of 4 mg/kg (Shear et al., 2002) delivered subcutaneously to the dorsal rump. Injections were subsequently given once daily for five days following injury (Shear et al., 2002). To allow clearance of progesterone and metabolites, the injection schedule ended more than 24 hrs before behavioral testing (Lin et al., 1972).

### Elevated Plus Maze

Ten days post-surgery, male and female rats were evaluated for generalized anxiety-like behavior using an elevated-plus maze (EPM) as described previously (Meyer et al., 2012, Fox et al., 2016, Scholl et al., 2019). Rats were placed in the center of the maze facing the closed arms and allowed to explore freely for 5 min. Time spent in the open arms and total distance moved in the maze were measured using Ethovision XT V5.1 (Noldus Information Technology, Leesburg, VA USA).

### Novel Object Recognition

Preliminary findings from the contextual fear experiments raised the possibility that progesterone was affecting learning or memory exclusively in male rats. To examine this further, subsequent cohorts of male rats were tested for short term memory prior to contextual fear conditioning. Short-term memory was examined using a novel object recognition (NOR) task, 24 hr after EPM testing with a 30 min delay (Fig. 1). To habituate to the testing arena, rats were allowed to freely explore the open field apparatus (98 L x 70 W cm) for 1 hr. The following day, rats were again placed into the open field for 5 min, then removed briefly to the home cage. Two identical objects (plastic bone or plastic golf ball) were placed at opposite ends of the arena and fixed in place. Rats were reintroduced and allowed to familiarize themselves with the objects for 10 min, and then returned to their home cage for the 30 min delay. The final trial used one familiar and one novel object in the arena (ball + bone, or bone + ball), and subjects explored the objects freely for 5 min. Time spent with each object was measured using EthovisionXT 8 software (Noldus Information Technologies), with the criterion that the nose of the rat was within 2 cm of the object. Discrimination index was calculated via the formula (Time Novel – Time Familiar) / (Time Novel + Time Familiar).

### Contextual Fear Conditioning

Assessment for contextual fear conditioning and extinction began 13 days after surgery (Figure 1). Testing occurred over the course of 4 days, as described in Meyer et al. (2012) and Davies et al. (2016). Briefly, on acquisition day, rats were placed in a foot shock chamber and were allowed to explore freely for 2 min. A total of 10 electric shocks (0.75 mA, 2 sec duration) was then delivered at intervals of 74 sec through the testing chamber floor (Meyer et al., 2012), with shock timing controlled by Ethovision 3.1 (Noldus Information Technologies). Rats then remained in the chamber for an additional 2 min with no shocks given before being removed. For test sessions 1–3, rats were placed each day for 8 min in the same chamber as on acquisition day, with no shocks delivered, to determine extent of contextual fear learning and extinction of fear conditioning. Video footage was scored later for freezing behavior using Observer XT10 (Noldus Information Technologies), by an observer blinded to treatment. Freezing behavior was defined as total immobility, except for those minor movements necessary for breathing (Forster et al. 2006).

One day following the final testing session, rats were anesthetized with sodium pentobarbital (100 mg/kg, IP) then perfused transcardially with phosphate-buffered saline (PBS, pH 7.4; room temperature), followed by 4% paraformaldehyde (pH 7.4, 4 °C). Brains and skulls were examined for evidence of hematoma, skull cracks or contusions, with brains retained for future study.

### Data Analysis

Separate t-tests compared social defeat scores between males and females, and time spent under anesthesia between control and defeat/mild TBI animals. Righting reflex times after surgery were compared using a two-way ANOVA (injury x sex) with subsequent multiple pairwise comparisons performed using Student-Newman-Keuls (SNK) tests.

To test the possible role of estrous cycle on behavioral outcomes, females were classified for each analysis by their estrous phase either on the day of mild TBI induction, or on the day of exposure to either the EPM or contextual fear conditioning (acquisition day). This allowed assessment of whether hormonal status interacts with mild TBI at the time of injury to drive behavioral changes, or alternatively, if it is the subsequent actions of ovarian steroids on the damaged neural landscape that is more important in promoting distinct phenotypes among estrous phases. Behavioral data were subjected to separate Grubbs’ tests for statistical outliers, resulting in removal of 5 out of 215 total data points for EPM data, 1 out of 40 for NOR data, and 78 out of 627 for fear conditioning data.

Separate three-way ANOVA (treatment x drug x sex/phase) were used to analyze time spent in the open arms of the EPM, as well as distance traveled in the maze, with interactions followed up by two-way ANOVA and SNK tests. Discrimination indices for NOR were analyzed by two-way ANOVA (treatment x drug) and post-hoc SNK tests. Total freezing time during contextual fear conditioning was analyzed across testing sessions separately within each sex/estrous cycle phase using 3-way mixed design ANOVA (treatment x drug x repeated measure of session), adjusted for non-sphericity using a Greenhouse-Geisser correction. Within each group, effects of session were followed by one-way repeated measures ANOVA to determine differences in freezing across sessions, with interactions followed by either 2-way mixed design ANOVA (repeated measure of session) or 2-way ANOVA (treatment x drug). Separate 3-way ANOVA (treatment x drug x sex/estrous cycle phase) were performed within each testing session to determine the effect of mild TBI and progesterone treatment on freezing behavior according to sex/ estrous phase. Multiple pairwise comparisons were carried out using Student-Newman-Keuls (SNK) tests. The significance level was set at P < 0.05 throughout, and all analyses were performed using either SigmaStat v13.0, or SPSS Version 24 (IBM Analytics).

## Results

### Social Defeat and Mild TBI Parameters

Social defeat scores did not differ between males (2.27 ± 0.13, mean score ± SE) and females (2.40 ± 0.12; t_147_ = -0.73, P = 0.466). Righting reflex time, a measure of the presence of mild TBI (Henninger et al., 2005, Meyer et al., 2012, Fox et al., 2016), was significantly greater in rats that received an impact (3.67 ± 0.06 min, mean ± SE) than those that received a sham surgery (2.38 ± 0.06 min; F_1, 275_ = 221.82; P < 0.001), with no difference between males and females (F_1,271_ = 0.05, P = 0.827), and no significant interaction between sex and injury (F_1,271_ < 0.001, P = 0.975). Rats receiving a mild TBI spent an average of 1.3 min longer under anesthesia than sham animals (t_276_ = 4.30, P < 0.001).

### Anxiety-like Behavior

#### Female estrous phase at time of injury

When females were classed by estrous phase at time of injury, 3-way ANOVA revealed only a main effect of sex/cycle phase on time spent in open arms of the EPM (F_4, 209_ = 8.96; P < 0.001), such that all females showed less anxiety-like behavior than males (SNK, P < 0.03 for all estrous phases; Table 1). Similarly, there was an effect of sex/cycle phase on distance moved (F_4, 209_ = 20.41; P < 0.001), with females overall traveling greater distances in the maze than males (SNK, P < 0.003 for all estrous phases; Table 2). There was no main effect of mild TBI on either open arm time (F_1, 209_ = 1.43; P = 0.23) or distance moved (F_1, 209_ = 2.18; P = 0.14) when females were grouped according to phase at injury, and no main effect of progesterone treatment on either variable (open arm time, F_1, 209_ = 0.03; P 0.87; distance moved, F_1, 209_ = 1.17; P = 0.28), and no interactions.

**Table 1.** Time spent in open arms (s) of the elevated plus maze when females were categorized by estrous phase at injury. Data are expressed as Mean + SEM. * = Difference from male within each treatment group (p < 0.05)

| Phase/Sex | Sham<br>Vehicle | Sham<br>Progesterone | Mild TBI<br>Vehicle | Mild TBI<br>Progesterone |
| --- | --- | --- | --- | --- |
| Male | 89.66 ± 13.41 | 84.67 ± 20.6 | 82.38 ± 10.43 | 61.07 ± 19.03 |
| Estrus | 118.49 ± 12.84* | 139.35 ± 6.48* | 125.20 ± 8.00* | 103.04 ± 17.87* |
| Metestrus | 117.15 ± 9.52* | 119.43 ± 11.13* | 133.50 ± 11.07* | 137.54 ± 4.42* |
| Diestrus | 128.23 ± 7.11* | 126.68 ± 8.48* | 120.48 ± 6.48* | 114.11 ± 9.34* |
| Proestrus | 130.67 ± 11.05* | 128.29 ± 6.89* | 111.19 ± 9.56* | 114.12 ± 16.45* |

**Table 2.** Distance moved (cm) in the elevated plus maze when females were categorized by estrous phase at injury. Data are expressed as Mean + SEM. * = Difference from male within treatment group (p < 0.05)

| Sex/phase | Sham<br>Vehicle | Sham<br>Progesterone | Mild TBI<br>Vehicle | Mild TBI<br>Progesterone |
| --- | --- | --- | --- | --- |
| Male | 2599.33 ± 73.59 | 2453.16 ± 114.4 | 2552.36 ± 78.83 | 2106.11 ± 123.31 |
| Estrus | 2631.80 ± 105.74* | 2734.65 ± 97.98* | 2840.8 ± 67.77* | 2736.29 ± 77.56* |
| Metestrus | 2805.6 ± 124.93* | 2830.9 ± 130.29* | 3033.7 ± 154.8* | 3179.39 ± 88.3* |
| Diestrus | 2937.28 ± 92.09* | 2920.63 ± 76.63* | 3076.1 ± 98.74* | 2834.18 ± 80.96* |
| Proestrus | 2816.97 ± 96.51* | 2824.86 ± 74.11* | 2788.1 ± 68.72* | 2876.6 ± 107.36* |

#### Female estrous phase at time of EPM testing

In contrast, when females were grouped according to estrous phase on the day of EPM testing, there were main effects of both mild TBI (F_1, 190_ = 7.18; P < 0.01) and sex/cycle phase (F_1, 190_ = 23.05; P < 0.001) on anxiety-like behavior, but no effect of progesterone treatment (F_1,190_ = 0.33; P = 0.57). There was also an interaction between mild TBI and sex/cycle phase (F_4,190_ = 4.40; P = 0.002). Males with mild TBI spent less time in the open arms of the maze than did sham males (SNK, P < 0.001; Fig. 2A). This same pattern was seen in diestrus females with mild TBI, which spent less time in the open arms than their uninjured controls (SNK, P = 0.025; Fig. 2A). However, no differences were seen in the time spent in open arms between mild TBI and sham females that were in either metestrus (SNK, P = 0.54), proestrus (SNK, P = 0.75) or estrus phase (SNK, P = 0.97) on the day of testing (Fig. 2A). Females as a whole also spent more time in open arms than males regardless of either injury or drug treatment (SNK, P < 0.02 for all estrous phases, Fig. 2A). Similarly, there was a main effect of sex/cycle phase (F_4,190_ = 16.48; P < 0.001) for distance traveled in the maze when females were classed by estrous phase at testing, but no effect of either mild TBI (F_1,190_ = 0.17; P = 0.68) or progesterone treatment (F_4,190_ = 0.33; P = 0.57), and no interactions, such that all females traveled greater distances in the maze than did males (SNK, P < 0.001 for all estrus phases; Fig. 2B). Subsequent linear regression analysis performed showed a weak correlation between distance moved by all females and time spent in open arms (r^2^ = 0.06; P = 0.002; Fig. 2C).

**Figure 2.**
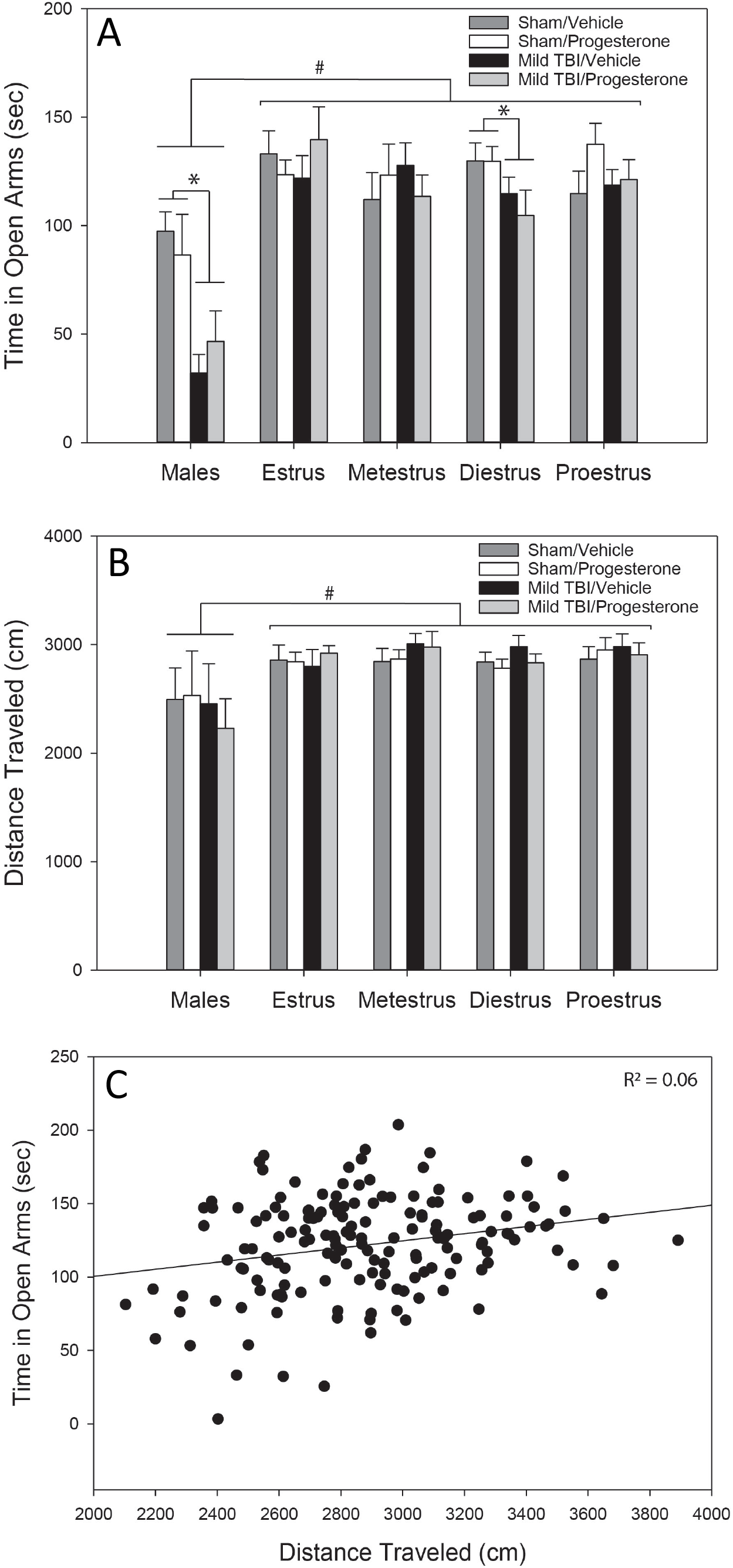
Anxiety-like Behavior in the Elevated Plus Maze. (**A**) Males (n = 10-14/group) and females in diestrus (n = 12-18/group) on the day of testing spend less time in the open arms of the maze following mild traumatic brain injury (Mild TBI) compared to sham controls in the same phase group (*P < 0.05) independent of progesterone treatment. Females spend more time in the open arms overall when compared to males (^#^P < 0.05). Estrus, metestrus and proestrus females show no differences among groups (n = 4-11 per group). (**B**) Females travel greater distances in the maze than males independent of injury or progesterone treatment (^#^P < 0.05). (**C**) A weak correlation is present between time spent in open arms and distance traveled in females (R^2^ = 0.06).

### Novel Object Recognition

No main effect of injury was evident in NOR, with control male rats discriminating between novel and familiar objects similarly to those with mild TBI (F_1, 35_ = 0.004; P = 0.947, Fig. 3). However, an effect of drug was observed (F_1, 35_ = 5.36; P = 0.027), in that all males treated with progesterone showed a reduced discrimination index compared to males treated with vehicle, independent of mild TBI status (Fig. 3).

**Figure 3.**
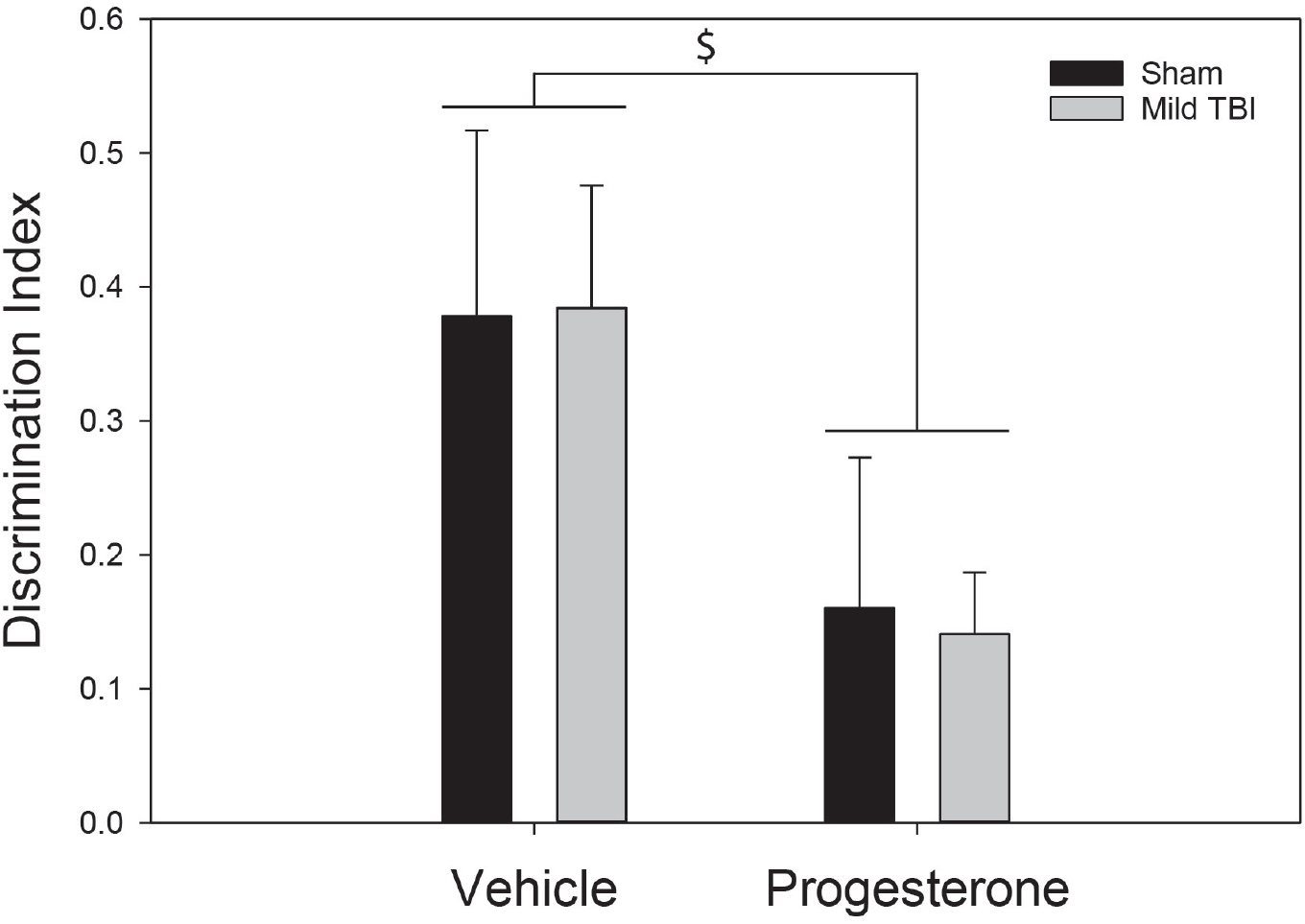
Novel Object Recognition in Male Rats. Males receiving progesterone treatment showed impaired ability to recognize a novel object compared to vehicle-treated rats (^$^P < 0.05). No difference in novel object recognition was seen between sham and mild traumatic brain injury (Mild TBI) groups treated with either vehicle or progesterone (n = 9-11 per group).

### Contextual Fear Conditioning

#### Comparing fear behavior across sessions within each sex/cycle phase

*Males* – There was no main effect of mild TBI on conditioned fear in male rats (F_1,45_ = 0.087; P = 0.77; Fig. 4), but there was an effect of drug (F_1,45_ = 15.55; P < 0.001) and an interaction between drug and session (F_1.1, 49.67_ = 7.58; P = 0.007). Both progesterone and vehicle treated males showed heightened freezing in session 1 that was extinguished by session 2 (SNK P < 0.001; Fig. 4). However, progesterone-treated males exhibited greater freezing than their vehicle-treated counterparts during sessions 1 and 2 (SNK P < 0.026; Fig. 4), with both groups exhibiting equivalent freezing in session 3 (SNK P > 0.11; Fig. 4).

**Figure 4.**
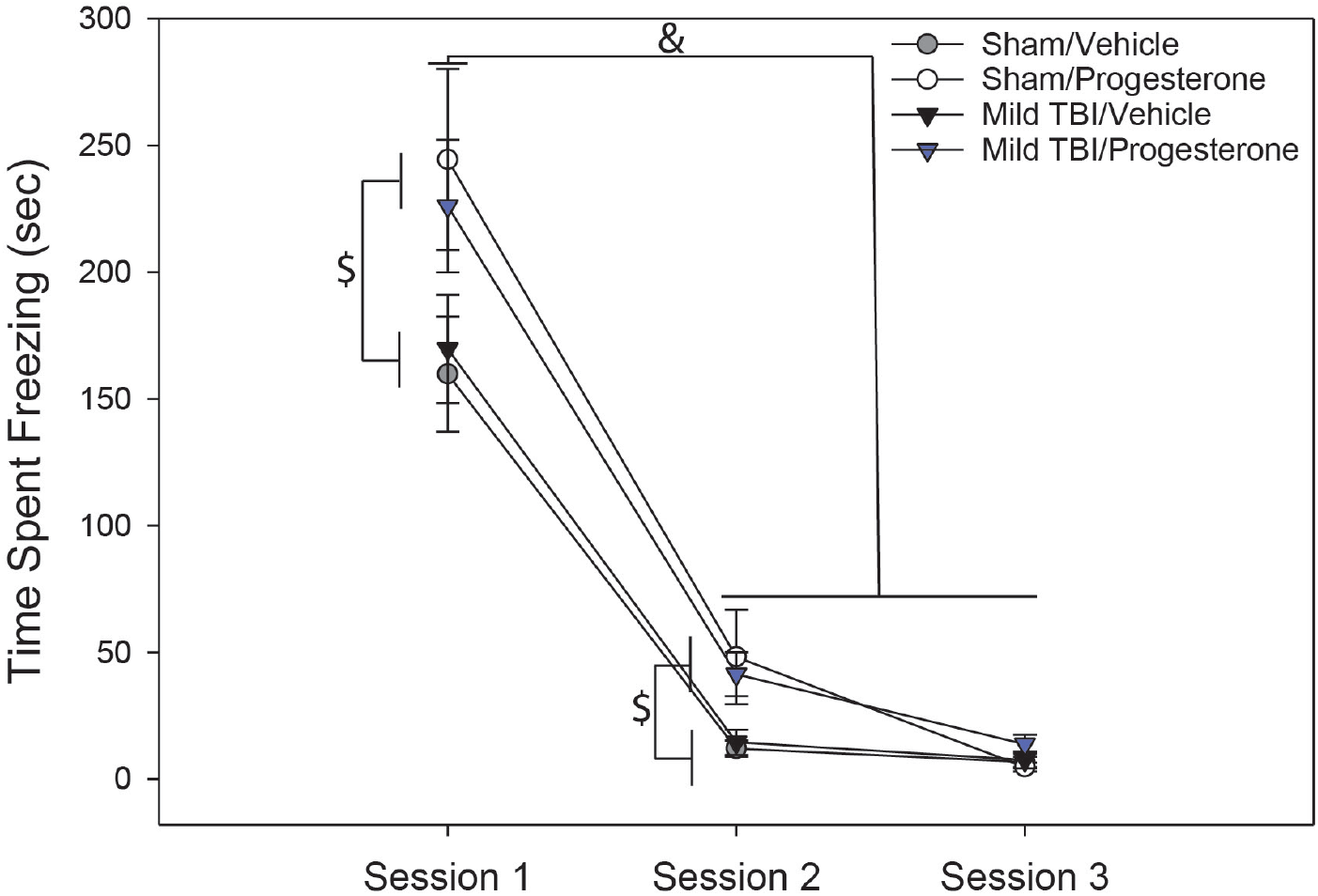
Contextual Fear Conditioning in Male Rats. Progesterone treatment enhanced conditioned freezing behavior during sessions 1 and 2 of testing (^$^P < 0.05) compared to vehicle controls. There was no effect of mild traumatic brain injury (Mild TBI), and all rats showed less conditioned freezing in sessions 2 and 3 compared to session 1 (^&^P < 0.05, n = 11-14 per group).

#### Female phase at time of injury

For female rats in estrus at time of injury, there was no effect of mild TBI on conditioned freezing (F_1,15_ = 0.3; P = 0.59). However, there was an effect of session (F_1.04,15.67_ = 25.8; P < 0.001), an effect of drug (F_1,15_ = 11.19; P = 0.004), and a drug x session interaction (F_1.04,15.67_ = 12.82; P = 0.002). During session 1, vehicle-treated females that were in estrus at time of injury showed higher levels of freezing than their progesterone-treated counterparts (SNK P < 0.001; Table 3). However, this difference according to drug treatment was not evident in sessions 2 and 3 (SNK P > 0.93; Table 3). Vehicle-treated estrus females also showed reductions in conditioned freezing from session 1 to session 2 (SNK P < 0.001; Table 3), while there was no difference in freezing across sessions for all progesterone-treated females that were in estrus at time of injury (SNK P > 0.18; Table 3).

**Table 3.** Time spent freezing (s) during conditioned fear recall when females were categorized by estrous phase at injury. Data are expressed as Mean + SEM. * = Difference from male (P < 0.05) within treatment group, # = Difference from vehicle within sex/phase (P < 0.05), *δ* = Difference from session 1 within treatment group (P < 0.001)

| Sex/phase | Treatment | Drug | N | Session 1 | Session 2 | Session 3 |
| --- | --- | --- | --- | --- | --- | --- |
| Male | Sham | Vehicle | 12 | 160.0±28.6 | 8.76±3.0$^{\delta}$ | 5.7±2.53$^{\delta}$ |
| | | Progesterone | 11 | 258.0±40.7$^{\#}$ | 38.0±15.4$^{\delta}$ | 5.71±2.7$^{\delta}$ |
| | Mild TBI | Vehicle | 12 | 162.0±33.1 | 8.93±2.28$^{\delta}$ | 3.41±1.53$^{\delta}$ |
| | | Progesterone | 14 | 271.0±26.3$^{\#}$ | 39.8±9.83$^{\delta}$ | 10.3±3.46$^{\delta}$ |
| Estrus | Sham | Vehicle | 4 | 140±47.4 | 1.43±0.82$^{\delta}$ | 0.27±0.12$^{\delta}$ |
| | | Progesterone | 6 | 23.3±13.5$^{* \#}$ | 0.7±0.6$^{*}$ | 0.0±0.0 |
| | Mild TBI | Vehicle | 3 | 150±74 | 14.9±7.7$^{\delta}$ | 0.69±0.12$^{\delta}$ |
| | | Progesterone | 6 | 30.3±15.7$^{* \#}$ | 8.63±7.6$^{*}$ | 1.13±1.1 |
| Metestrus | Sham | Vehicle | 10 | 85.3±38.8$^{*}$ | 13.5±7.8$^{\delta}$ | 3.74±1.5$^{\delta}$ |
| | | Progesterone | 8 | 132±50.6$^{*}$ | 8.24±6.2$^{* \delta}$ | 23.6±23.4$^{\delta}$ |
| | Mild TBI | Vehicle | 7 | 128±49.4$^{*}$ | 4.37±2.5$^{\delta}$ | 0.1±0.08$^{\delta}$ |
| | | Progesterone | 6 | 26.3±14.4$^{*}$ | 3.0±2.2$^{* \delta}$ | 0.24±0.16$^{\delta}$ |
| Diestrus | Sham | Vehicle | 18 | 90.0±25.5$^{*}$ | 2.05±0.84$^{\delta}$ | 0.77±0.29$^{\delta}$ |
| | | Progesterone | 12 | 71.3±27.2$^{*}$ | 0.46±0.27$^{* \delta}$ | 1.23±0.36$^{\delta}$ |
| | Mild TBI | Vehicle | 18 | 68.7±26.1$^{*}$ | 5.59±2.65$^{\delta}$ | 0.63±0.35$^{\delta}$ |
| | | Progesterone | 14 | 61.6±20.8$^{*}$ | 1.49±0.99$^{* \delta}$ | 1.73±1.3$^{\delta}$ |
| Proestrus | Sham | Vehicle | 8 | 74.9±36.7$^{*}$ | 2.13±1.37$^{\delta}$ | 0.40±0.35$^{\delta}$ |
| | | Progesterone | 11 | 88.1±25.3$^{*}$ | 4.67±2.6$^{* \delta}$ | 0.54±0.31$^{\delta}$ |
| | Mild TBI | Vehicle | 10 | 36.1±13.7$^{*}$ | 2.0±1.64$^{* \delta}$ | 0.34±0.22$^{\delta}$ |
| | | Progesterone | 10 | 111±36.5$^{*}$ | 0.76±0.44$^{* \delta}$ | 2.41±1.42$^{\delta}$ |

Separate 3-way repeated measures ANOVA within all other phases (proestrus, metestrus, diestrus), as classed at time of injury, revealed no effect of either mild TBI or progesterone on freezing behavior across sessions in any group (P > 0.07), and no interactions between these two factors. There was a main effect of session for each phase (proestrus: F_1.01,35.22_ = 27.99; P < 0.001; estrus: F_1.03,17.44_ = 18.34; P < 0.001; metestrus: F_2,54_ = 18.67; P < 0.001; diestrus: F_2,116_ = 31.36; P < 0.001), with all females showing reductions in conditioned freezing from sessions 1 to 2 (SNK P < 0.001; Table 3).

#### Female phase at fear acquisition

In contrast, for female rats that were in estrus on the day of fear acquisition (shock + context pairing), there were main effects of session (F_1,20_ = 14.73; P < 0.001) and mild TBI (F_1,10_ = 6.91; P = 0.025), but no effect of progesterone (F_1,10_ = 0.59; P = 0.46). There was also an interaction (F_1,62_ = 119.19; P < 0.001) between mild TBI and session for this group, such that estrus females with mild TBI exhibited less freezing behavior than sham females, but only during session 1 (SNK, P = 0.025; Fig. 5D). Sham females in estrus also showed a decrease in conditioned freezing from session 1 to session 2 (SNK P < 0.001; Fig. 5D), while time spent freezing did not change across sessions for mild TBI females in estrus (SNK P = 0.18; Fig. 5D).

**Figure 5.**
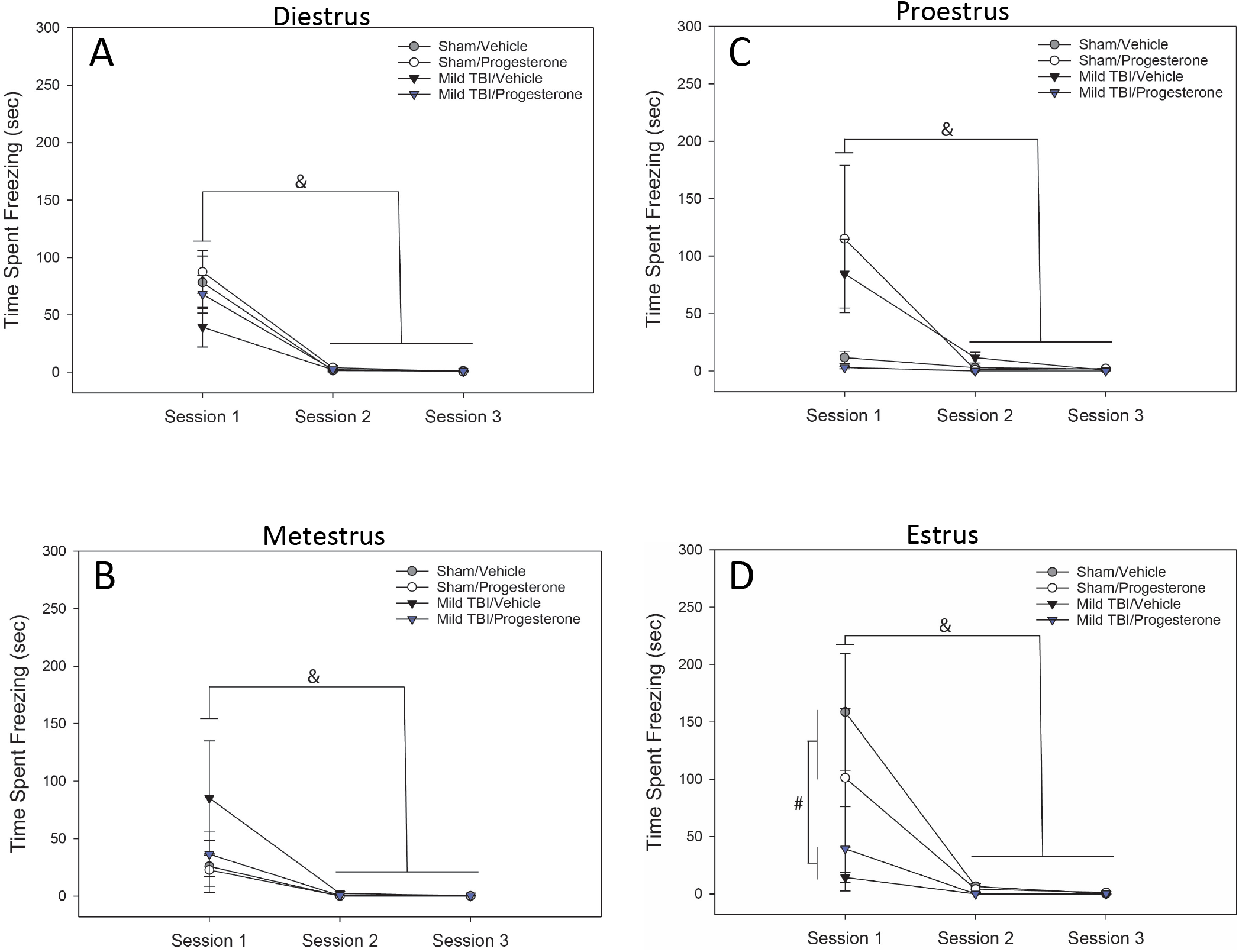
Contextual Fear Conditioning in Female Rats. (**A**) Females in diestrus phase on the day of fear acquisition go on to extinguish conditioned fear in sessions 2 and 3 (^&^P < 0.05), but otherwise show no response to either mild traumatic brain injury (Mild TBI) or progesterone treatment (n = 15-19 per group). Similar results were seen in (**B**) metestrus (n = 3-10 per group) and (**C**) proestrus females (n = 3-11 per group). (**D**) Females in estrus on the day of fear acquisition also demonstrated an effect of mild TBI (^#^P < 0.05) in session 1 (n = 3-4 per group).

Females in proestrus at acquisition showed an interaction among injury, drug and session (F_1.02,26.47_ = 5.29; P = 0.029), along with a main effect of session (F_1.02,26.47_ = 9.28; P = 0.005). Subsequent two-way ANOVA within each session revealed an interaction between injury and drug during session 1 (F_1,10_ = 6.91; P < 0.025), which was not present during either session 2 (F_1,22_ = 0.92; P = 0.35) or 3 (F_1,22_ = 0.23; P = 0.64). During session 1, vehicle-treated proestrus females with mild TBI exhibited greater conditioned freezing than vehicle-treated sham females (SNK P = 0.03; Fig. 5C). There was a trend for progesterone treatment to ameliorate the increased freezing in females with mild TBI (Fig. 5C), but this did not reach statistical significance (SNK P = 0.076). Conversely, there was a non-significant trend for progesterone to increase freezing in sham females in proestrus at time of fear conditioning (SNK P = 0.058; Fig. 5C). Decreases in conditioned freezing from session 1 to sessions 2 and 3 were seen in each group of proestrus females (SNK P < 0.049; Fig. 5C), with the exception of the Sham+Vehicle rats (SNK P = 0.12; Fig. 5C).

There was no effect of either mild TBI (P > 0.07) or progesterone (P > 0.45) on freezing in females that were in metestrus or diestrus at time of fear acquisition, and no interactions between these factors. Females in both phases did show a main effect of session (metestrus: F_1, 25.02_ = 5.66; P = 0.025; diestrus: F_1.01, 61.43_ = 48.66; P < 0.001), with conditioned fear being highest during session 1 and extinguishing by sessions 2 and 3 (SNK P < 0.001; Figs 5A and B).

#### Comparing fear behavior among sex/estrous cycle phase within each session

Separate 3-way ANOVA (treatment x drug x sex/cycle phase) revealed an interaction between drug and sex/cycle phase on freezing during session 1, which was evident when females were classed by estrous phase at either time of injury (F_4,182_ = 4.15; P = 0.003) or at time of fear acquisition (F_4,164_ = 3.1; P = 0.017). Vehicle-treated females that were in estrus at time of injury exhibited equivalent freezing to vehicle-treated males (SNK P = 0.41; Table 3). In contrast, there was a trend for freezing to be higher in vehicle-treated males when compared to vehicle-treated females that were in estrus at time of fear acquisition, although this did not reach statistical significance (P = 0.054; Table 4). With the exception of females in estrus, all vehicle-treated male rats exhibited greater conditioned fear than all other vehicle-treated females during session 1, regardless of when estrous phase was classed (SNK P < 0.027; Tables 3 and 4). Progesterone treatment caused males to display higher freezing in session 1 than all progesterone-treated females, again irrespective of when estrous phase was classed (SNK P < 0.001; Tables 3 and 4).

**Table 4.**
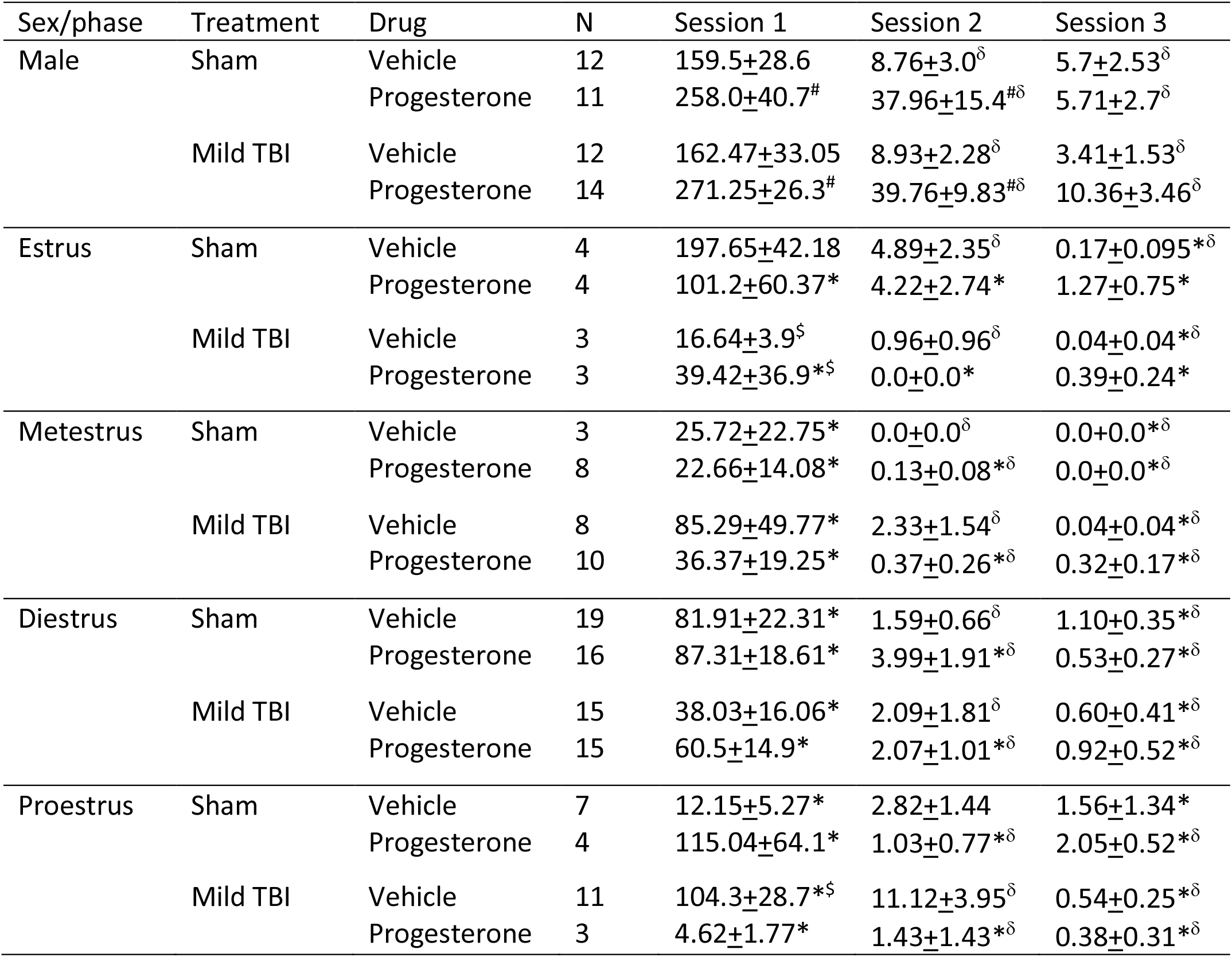
Time spent freezing (s) during conditioned fear recall when females were categorized by estrous phase at fear acquisition. Data are expressed as Mean + SEM. * = Difference from male (P < 0.05) within treatment group, # = Difference from vehicle within sex/phase (P < 0.05), $ = Difference from sham within treatment group (P = 0.025); *δ* = Difference from session 1 within treatment group (P < 0.001).

| Sex/phase | Treatment | Drug | N | Session 1 | Session 2 | Session 3 |
| --- | --- | --- | --- | --- | --- | --- |
| Male | Sham | Vehicle | 12 | 159.5±28.6 | 8.76±3.0δ | 5.7±2.53δ |
| | | Progesterone | 11 | 258.0±40.7<sup>#</sup> | 37.96±15.4<sup>#$\delta$</sup> | 5.71±2.7δ |
| | Mild TBI | Vehicle | 12 | 162.47±33.05 | 8.93±2.28δ | 3.41±1.53δ |
| | | Progesterone | 14 | 271.25±26.3<sup>#</sup> | 39.76±9.83<sup>#$\delta$</sup> | 10.36±3.46δ |
| Estrus | Sham | Vehicle | 4 | 197.65±42.18 | 4.89±2.35δ | 0.17±0.095<sup>*δ</sup> |
| | | Progesterone | 4 | 101.2±60.37<sup>*</sup> | 4.22±2.74<sup>*</sup> | 1.27±0.75<sup>*</sup> |
| | Mild TBI | Vehicle | 3 | 16.64±3.9<sup>\$</sup> | 0.96±0.96δ | 0.04±0.04<sup>*δ</sup> |
| | | Progesterone | 3 | 39.42±36.9<sup>*\$</sup> | 0.0±0.0<sup>*</sup> | 0.39±0.24<sup>*</sup> |
| Metestrus | Sham | Vehicle | 3 | 25.72±22.75<sup>*</sup> | 0.0±0.0δ | 0.0±0.0<sup>*δ</sup> |
| | | Progesterone | 8 | 22.66±14.08<sup>*</sup> | 0.13±0.08<sup>*δ</sup> | 0.0±0.0<sup>*δ</sup> |
| | Mild TBI | Vehicle | 8 | 85.29±49.77<sup>*</sup> | 2.33±1.54δ | 0.04±0.04<sup>*δ</sup> |
| | | Progesterone | 10 | 36.37±19.25<sup>*</sup> | 0.37±0.26<sup>*δ</sup> | 0.32±0.17<sup>*δ</sup> |
| Diestrus | Sham | Vehicle | 19 | 81.91±22.31<sup>*</sup> | 1.59±0.66δ | 1.10±0.35<sup>*δ</sup> |
| | | Progesterone | 16 | 87.31±18.61<sup>*</sup> | 3.99±1.91<sup>*δ</sup> | 0.53±0.27<sup>*δ</sup> |
| | Mild TBI | Vehicle | 15 | 38.03±16.06<sup>*</sup> | 2.09±1.81δ | 0.60±0.41<sup>*δ</sup> |
| | | Progesterone | 15 | 60.5±14.9<sup>*</sup> | 2.07±1.01<sup>*δ</sup> | 0.92±0.52<sup>*δ</sup> |
| Proestrus | Sham | Vehicle | 7 | 12.15±5.27<sup>*</sup> | 2.82±1.44 | 1.56±1.34<sup>*</sup> |
| | | Progesterone | 4 | 115.04±64.1<sup>*</sup> | 1.03±0.77<sup>*δ</sup> | 2.05±0.52<sup>*δ</sup> |
| | Mild TBI | Vehicle | 11 | 104.3±28.7<sup>*\$</sup> | 11.12±3.95δ | 0.54±0.25<sup>*δ</sup> |
| | | Progesterone | 3 | 4.62±1.77<sup>*</sup> | 1.43±1.43<sup>*δ</sup> | 0.38±0.31<sup>*δ</sup> |

A drug x sex/cycle phase interaction was also present for freezing during session 2 (phase at injury: F_4,201_ = 7.1; P < 0.001; phase at fear acquisition: F_4,164_ = 6.93; P < 0.001). However, sex differences in fear behavior during session 2 were restricted to progesterone-treated rats, with males freezing more than females in any estrous phase, whether this was categorized at time of injury or at fear acquisition (SNK P < 0.001; Tables 3 and 4), while there were no differences among vehicle-treated males and females (P > 0.55; Tables 3 and 4). By session 3, there was only an effect of phase/sex when females were classed by estrous phase at time of fear acquisition (F_4,164_ = 10.12; P < 0.001), with all males showing greater levels of freezing than all females. There were no differences seen among female cycle phases in time spent freezing during any session (SNK P > 0.32 for all estrous phase comparisons within each session for phase at injury and phase at acquisition; Tables 3 and 4).

## Discussion

This study sought to determine potential benefits of progesterone treatment on preventing anxiety- and PTSD-like behaviors following mild TBI in male and naturally cycling female rats. Mild TBI and progesterone treatment differentially affected behavior depending on both sex and estrous cycle phase, and these effects were specific to the behavior being assessed. Mild TBI increased anxiety-like behavior in male rats, but alterations to male fear learning were solely a response to progesterone treatment. Conversely, mild TBI decreased fear conditioning in females, but only for those in estrus at the time of fear acquisition, and no effect of injury on anxiety was seen in this group. In contrast, mild TBI increased conditioned fear in females that were in proestrus at time of acquisition, but only in those treated with vehicle. Interestingly, effects of mild TBI on anxiety and fear conditioning in females were not evident when grouped by estrous phase at time of injury, suggesting differential outcomes of mild TBI in females are dependent on actions of ovarian steroid fluctuations subsequent to neural damage, and not when the injury is initially sustained. These findings highlight the importance of considering both sex and female reproductive cycle phase when investigating how hormonal state and/or treatment may either promote or protect against the negative outcomes of mild TBI.

Previous work in male rats has shown that the closed-head mild TBI with a social stressor reliably produces anxiety- and PTSD-like behaviors (Meyer et al., 2012; Davies et al., 2016; Fox et al., 2016). To better understand sex differences in human responses to mild TBI, this study included naturally cycling female rats. As the first study to use a combined social stress and weight drop model of concussive injury in female rats, we demonstrated that both males and females received comparable aggression during social defeat exposure. Similarly, we confirmed that there were no sex differences observed in the time to obtain a righting reflex following mild TBI. Increased latency to exhibit the righting reflex in male rats is characteristic of mild TBI (Meyer et al., 2012, Davies et al., 2016, Fox et al., 2016), and our results suggest that this is also occurs in females. Together, the similarities in both the stress and mild TBI-inducing components of the model validates its use in both sexes.

Male rats demonstrated greater anxiety-like behavior following mild TBI, replicating our previous findings (Davies et al., 2016; Fox et al., 2016), and consistent with increased anxiety reported in human males who have suffered concussive injury (Moore et al., 2006, Hoge et al., 2008). Previous work has shown that the combination of mild TBI and increased circulating glucocorticoids (as induced by social defeat at the time of injury) produces greater anxiety than either mild TBI or social stress alone (Davies et al., 2016; Fox et al., 2016), suggesting an additive effect of injury and stress on negative outcomes. In female rats, only those in diestrus phase at the time of testing showed increased anxiety-like behavior following mild TBI. Diestrus is the longest phase of the rat reproductive cycle, lasting between 24 and 48 hours, and during this time endogenous ovarian sex steroids are at their lowest blood levels (Schumacher et al., 2014). Since male rats have comparably low levels of circulating estradiol and progesterone, it may be that the natural current or recent elevation in these hormones masked the anxious phenotype in estrus, proestrus and metestrus rats that had received mild TBI. Comparing anxiety-like behavior following mild TBI in ovariectomized females and those receiving exogenous steroid replacement just prior to testing would be valuable in elucidating the contribution of endogenous estrogen and progesterone to the anxiogenic effects of mild TBI. Similarly, assessing the effect of proestrus-comparable doses of ovarian sex steroids given to males on the day of EPM testing could provide even further understanding.

Progesterone treatment at 4 mg/kg for five days after injury failed to prevent emergence of anxiety-like behavior following mild TBI in either sex, or in any estrous phase. It has been suggested that the beneficial effects of progesterone following mechanical neuronal insult are optimal when combined with other treatments, including enriched environment (Nudi et al., 2015) or vitamin D treatment (Tang et al., 2015), and that monotherapy may not be adequate. The 4 mg/kg of progesterone used here may not be sufficient to prevent the anxiogenic effects of mild TBI, although this dose has been shown to prevent edema after brain injury in both sexes (Roof et al., 1992). The effects of much higher doses of progesterone in more severe cases of TBI have been explored in juvenile (4 weeks old) rats (8 mg/kg in females, Geddes et al., 2016, or even 16 mg/kg in males, Geddes et al., 2014), and these doses improved performance spatial navigation/spatial memory as shown in a Morris water maze task. However, 4 mg/kg for five days was effective in preventing TBI-induced cognitive deficits in spatial learning and attention in male rats (Shear et al., 2002). The detrimental effects of 4 mg/kg progesterone in enhancing fear conditioning and impairing novel object recognition in male rats in our study suggest that a higher concentration of progesterone would not be appropriate as a post-mild TBI anxiolytic treatment in this sex. However, a dose-response curve for progesterone’s actions on anxiety-like behavior may yield interesting insights, perhaps not following a linear curve, and could be examined in future work.

It is interesting that although the stage of the estrous cycle in female rats affected the outcome of mild TBI on anxiety-like behaviors, progesterone treatment itself did not have an impact. As in previous work, females traveled significantly further in the maze overall than males regardless of cycle stage (Scholl et al., 2019), though regression analysis showed only 6% of variation in time spent in the open arms could be accounted for by distance traveled in female rats. Female rats naturally ambulate more than males in the EPM (Scholl et al., 2019, Santos et al., 2014, Simpson et al., 2012, Devall et al., 2009, Johnston and File, 1991). Such differences have been attributed to ovarian sex steroids (Walf et al., 2008), although the current study does not show variation in locomotion in the EPM as a function of estrous cycle and associated ovarian steroid fluctuations. Overall, variance in distance traveled within the EPM is unlikely to explain the differential effects of mild TBI on anxiety-like behavior in male versus female rats seen here.

In contrast to anxiety-like behavior, contextual fear conditioning showed no influence of mild TBI in males but an effect of progesterone. Male rats with mild TBI did not demonstrate an expected elevation in fear conditioning or impairment in fear extinction, failing to replicate a prior study in which the concurrent-social stressor-with-injury model produced these responses (Davies et al., 2016). While the duration of freezing behavior following mild TBI was similar, sham animals showed enhanced fear conditioning relative to that reported by Davies et al. (2016), resulting in the mild TBI and sham groups being statistically indistinguishable. All male rats in this study conditioned to the shock-associated context and extinguished freezing behavior over the subsequent sessions. However, progesterone treatment produced a significant increase in freezing behavior compared to vehicle treatment in session 1, which persisted into session 2 (48 hours after acquisition). This suggests progesterone treatment not only enhanced fear conditioning, but may have also produced an impairment in fear extinction. This finding is particularly interesting considering that the final progesterone treatment occurred four days prior to fear conditioning testing, suggesting persistence of progesterone’s effects despite a reduction in circulating levels.

A possible explanation for this result is that repeated progesterone treatment may have enhanced learning processes in males, promoting a greater conditioned fear response. Progesterone delivery in prior studies improved cognitive deficits following more severe TBI in the Morris Water Maze, a task requiring intact learning ability for optimal performance (Si et al., 2013, Shear et al., 2002).

Progesterone treatment in these studies usually ended a day prior to behavioral testing (Shear et al., 2002). However, it is not known whether progesterone treatment that ceases four or more days prior to testing affects learning. To examine this, we conducted NOR testing with male rats using a 30 min delay, seeking specifically to assess hippocampal function (Han et al., 2014, Park et al., 2016) known to be important for contextual fear conditioning (Maren et al., 2013). Unexpectedly, progesterone treatment produced a significant impairment in the ability of male rats to discriminate a novel from a familiar object, even though the last injection of progesterone occurred four days prior to testing. This finding suggests that the enhanced fear conditioning response due to progesterone treatment in males is unlikely to be due to a general enhancement in learning.

It was also initially surprising that mild TBI did not affect NOR in male rats. However, studies that show mild TBI-induced impairment in NOR often conduct the test within a few days after injury (Baratz-Goldstein et al., 2016, Munyon et al., 2014). Consistent with the current study, studies conducting NOR testing more than seven days following mild TBI find no impairment following injury, or that a prior deficit improves with a similar lapse in time (Yang et al., 2013, Rachmany et al., 2013). Therefore, short-term memory deficits may be transient in male rodent models of mild TBI. It is worth mentioning that the current NOR testing utilized different objects in a fixed position in the testing arena, whereas alternative NOR paradigms involve moving objects around within the open field, employing components of hippocampal spatial learning in addition to object memory (Barker et al., 2007). Long-lasting deficits in the spatially-based Morris Water Maze as a result of mild TBI have been noted in male rats (Henninger et al., 2005) and improvements have been reported following progesterone administration after more severe TBI (Shear et al., 2002). Therefore, a spatial version of the NOR test may reveal an effect of mild TBI, and/or progesterone treatment.

It is not clear why five days of progesterone treatment after mild TBI would enhance fear conditioning while impairing fear extinction and hippocampal-based memory. One potential mechanism could be a result of negative feedback, where repeated progesterone treatment downregulated normal sex hormone production in males, resulting in atypically low levels of neuroprotective steroids (including progesterone) at the time of behavioral assessment. An acute loss of allopregnanolone, a predominant progesterone metabolite in the brain, may depress GABAergic signaling in a state dependent manner (Acca et al., 2017), leading to over-activity of brain regions governing fear expression such as the amygdala and hippocampus. It is also known that decreases in allopregnanolone in the brains of male rats are associated with enhanced fear conditioning (Zhang et al., 2016). Further studies that assess circulating and brain hormone levels over the course of progesterone treatment as along with GABAergic signaling following treatment will be useful in determining these mechanisms. Regardless of what potential mechanisms may be driving the downstream effects, this study demonstrates that progesterone treatment may not be the optimal approach for treating post-concussive anxiety and PTSD-like symptoms in males.

Female rats expressed less conditioned freezing overall than male rats, a finding supported by previous studies using contextual fear learning paradigms (Cossio et al., 2016, Daviu et al., 2014, Chang et al., 2009, Maren et al., 1994). In contrast to other studies reporting lower conditioned fear (contextual or cued) in proestrus females compared to other phases of the estrous cycle (Milad et al., 2009, Markus and Zecevic 1997), we observed no differences among females as a function of reproductive cycle stage. In addition, the majority of female rats exposed to mild TBI did not exhibit the expected increase in fear conditioning, with the exception of rats that were in proestrus at time of fear acquisition, but this was restricted to those treated with vehicle. There was also a trend for heightened freezing by injured proestrus females to be ameliorated by progesterone treatment, but this failed to achieve statistical significance (P = 0.076), possibly because of the low sample size in this particular group. In contrast, blunted conditioned freezing was observed in mild TBI females that were in estrus on the day of fear acquisition, which was independent of progesterone treatment. The variance in female fear conditioning data as a result of allowing females to cycle naturally throughout the experiment, and inclusion of both early- and late-estrus females in that group, suggest that follow-up studies are required to validate these findings. If replicated, reduced fear conditioning in the estrus phase suggests that a concussed female may be more protected from some PTSD-like symptoms if trauma exposure occurs shortly following ovulation before the metestrus phase begins. Conversely, prophylactic delivery of progesterone may mitigate negative effects of mild TBI when subsequent trauma is experienced prior to ovulation, but this also requires confirmation with further studies.

## Conclusions

Progesterone has shown great promise in early studies as a potential therapeutic agent following major head trauma (Roof et al., 1996, Shear et al., 2002, Sayeed and Stein 2009, Geddes et al., 2014). Our results show progesterone did not attenuate psychological symptoms of mild TBI in male or female rats. Furthermore, progesterone treatment of male rats impaired hippocampal-based memory and enhanced contextual fear conditioning. Based on these results, monotherapy with progesterone is unlikely to be a successful candidate as a treatment for post-concussive anxiety or PTSD in males. In female rats, future directions are less clear. Mild TBI paired with social stress had little effect on anxiety or fear conditioning in females, and effects observed were confined to specific phases of the reproductive cycle, and only when phase was classed at the time of behavioral testing. A more robust animal model of female concussive injury that better mimics the human state may need to be established to determine the full potential for progesterone to ameliorate the psychological effect of mild TBI in females. The psychological changes that occur in many recipients of mild TBI are an important facet of the condition, and our results reiterate that males and females might express these changes differently and may also respond differently to progesterone as a post-concussive treatment. In summary, it is clear that taking these differences into account may allow for an improved ability to individualize treatments for all who receive a concussive injury, regardless of sex or current reproductive phase.

## Abbreviations

(EPM): Elevated-plus maze
(NOR): Novel Object Recognition
(PTSD): Posttraumatic stress disorder
(Prog): Progesterone
(SNK): Student-Newman-Keuls
(TBI): Traumatic brain injury

## Acknowledgements

The authors would like to thank Nate J. Vinzant, Matthew A. Weber, Brenna Bray, Shaydel R. Engel and Riley T. Paulsen for their contributions to this work, Dr. Kenneth J. Renner for mentorship and manuscript preparation, as well as the Animal Behavioral Testing Core at the University of South Dakota (USD).

## Highlights

Sex differences evident in a rat model of PTSD and mild traumatic brain injury (mTBI)

Progesterone treatment does not attenuate behavioral outcomes of mTBI

Estrous cycle alters female response to anxiety-like behavior and fear conditioning

